# Divergent changes in PBN excitability in a mouse model of neuropathic pain

**DOI:** 10.1101/2023.10.11.561891

**Authors:** María L Torruella-Suárez, Benjamin Neugebauer, Krystal Flores-Felix, Asaf Keller, Yarimar Carrasquillo, Nathan Cramer

## Abstract

The transition from acute to chronic pain involves maladaptive plasticity in central nociceptive pathways. Growing evidence suggests that changes within the parabrachial nucleus (PBN), an important component of the spino–parabrachio–amygdaloid pain pathway, are key contributors to the development and maintenance of chronic pain. In animal models of chronic pain, PBN neurons become sensitive to normally innocuous stimuli and responses to noxious stimuli become amplified and more often produce after-discharges that outlast the stimulus. Using *ex vivo* slice electrophysiology and two mouse models of neuropathic pain, sciatic cuff and chronic constriction of the infraorbital nerve (CCI-ION), we find that changes in the firing properties of PBN neurons and a shift in inhibitory synaptic transmission may underlie this phenomenon. Compared to PBN neurons from shams, a larger proportion of PBN neurons from mice with a sciatic cuff were spontaneously active at rest, and these same neurons showed increased excitability relative to shams. In contrast, quiescent PBN neurons from cuff mice were less excitable than those from shams. Despite an increase in excitability in a subset of PBN neurons, the presence of after-discharges frequently observed *in vivo* were largely absent *ex vivo* in both injury models. However, GABA_B_-mediated presynaptic inhibition of GABAergic terminals is enhanced in PBN neurons after CCI-ION. These data suggest that the amplified activity of PBN neurons observed in rodent models of chronic pain arise through a combination of changes in firing properties and network excitability.

**Significance Statement:** Hyperactivity of neurons in the parabrachial nucleus (PBN) is causally linked to exaggerated pain behaviors in rodent models of chronic pain but the underlying mechanisms remain unknown. Using two mouse models of neuropathic pain, we show the intrinsic properties of PBN neurons are largely unaltered following injury. However, subsets of PBN neurons become more excitable and GABA_B_ receptor mediated suppression of inhibitory terminals is enhanced after injury. Thus, shifts in network excitability may be a leading factor in injury induced potentiation of PBN activity.

## Introduction

The parabrachial nucleus (PBN) is an important sensory processing and relay station for pain neurotransmission (Chiang et al., 2019; Palmiter, 2018). It is the main target of lamina I spinal nociceptive neurons (Barik et al., 2018; Bernard et al., 1989; Cechetto et al., 1985; Craig, 1995), and receives strong input from Tac1-expressing cells that transmit nociceptive information (Al-Khater and Todd, 2009; Chiang et al., 2020). Histological (Bellavance and Beitz, 1996; Bester et al., 2000, 1997; Chiang et al., 2020; Dutschmann and Herbert, 1997; Hermanson and Blomqvist, 1997, 1996; Jergova et al., 2008; Li et al., 2006; Qi et al., 2022), electrophysiological (Bernard et al., 1994; Bernard and Besson, 1990; Bourgeais et al., 2001; Uddin et al., 2018), and calcium imaging studies (Barik et al., 2021; Campos et al., 2018), largely in rodents, established that PBN neurons respond to noxious stimulation in both naïve and chronic pain conditions. PBN neurons become hyperresponsive in several models of chronic pain (Matsumoto et al., 1996; Raver et al., 2020; Sun et al., 2020; Uddin et al., 2018), and have increased spontaneous activity (Matsumoto et al., 1996; Raver et al., 2020). After discharges, where PBN neurons remain active long after the noxious stimulus has ended, are also more prevalent in rodents with chronic pain (Uddin et al., 2018). Suppressing this enhanced excitability also suppresses chronic pain behaviors (Raver et al., 2020), suggesting changes in PBN excitability are causally linked to changes in pain perception.

Shifts in PBN excitability may arise from a combination of factors, including changes in the intrinsic properties of the neurons and changes in the strength of afferent inputs. Among the many neurotransmitters and neuromodulators in the PBN (Cramer et al., 2021; Teuchmann et al., 2022), GABA has emerged as a crucial locus of injury-induced plasticity. The central nucleus of the amygdala (CeA) sends GABAergic inputs to the PBN which are attenuated in a model of chronic pain (Raver et al., 2020). Furthermore, optogenetic stimulation of this input can ameliorate pain-related behaviors (Hogri et al., 2022; Raver et al., 2020). The CeA is likely a source of tonic GABA to PBN in intact animals, as inhibition of part of this input is aversive even in the absence of injury (Torruella-Suárez et al., 2020). The PBN is under tonic suppression also by local GABAergic interneurons as optogenetic inhibition of this population can induce tactile hypersensitivity in otherwise pain-free animals (Sun et al., 2020). These studies position GABA as a key modulator of PBN activity in pain states.

While a wealth of literature has linked changes in PBN activity with chronic pain behaviors, the mechanisms driving these changes are largely unknown. In this study we examined whether PBN hyperexcitability described *in vivo* is due to changes in intrinsic excitability and GABAergic signaling. As the previously described histological studies largely found differences in Fos expression in the lateral subnucleus of the PBN (lPBN), (Bellavance and Beitz, 1996; Bester et al., 2000; Dutschmann and Herbert, 1997; Hermanson and Blomqvist, 1997, 1996; Jergova et al., 2008; Li et al., 2006), the lPBN receives dense spinal inputs (Choi et al., 2020), and lPBN outputs have been linked to nocifensive behaviors (Chiang et al., 2020), we focused our recordings on this region. We further characterize passive and active membrane properties as well as spontaneous and evoked repetitive firing of PBN neurons, both at baseline and following nerve injury using two rodent models of neuropathic pain. Lastly, we tested the hypothesis that GABAergic input onto PBN neurons is subject to injury-induced plasticity. Our combined results demonstrate that lateral PBN neurons become more excitable following peripheral nerve injury and further suggest that enhancement of GABA_B_-mediated presynaptic inhibition contributes to pain-related plasticity.

## Methods

### Subjects

All animal procedures were performed in accordance with the [Author University] animal care committee’s regulations. We used male and female C57BL/6J mice, but some experiments were performed only in male mice. These data are noted accordingly. All mice were bred in house or purchased from Jackson Laboratory and were 8 to 16 weeks old at the time of experimentation. Animals were group-housed prior to surgery in a reverse 12/12 light cycle with *ad libitum* food and water.

### Sciatic cuff surgery

The sciatic cuff model of neuropathic pain was originally designed by Benbouzid *et al* (2008). Male mice were anesthetized using isoflurane with an initial dose of 2% and a maintenance dose of 1% using a VetFlo anesthesia system (Kent Scientific, Torrington, CT, US). A hand warmer was used for warmth during the procedure and recovery. The right sciatic nerve was exposed, and a 2 mm piece of polyethylene tubing (0.38 mm ID/1.09 mm OD) was slid onto the nerve of cuff animals. The nerve was then returned to its original position and wound clips were used to close the surgical site. Sham animals underwent the same process without cuff placement. Animals were placed in a new, clean cage and allowed to recover. Following surgery, mice were single- or pair-housed with a littermate that had received the same surgical intervention (sham or cuff) in a cage with a perforated Plexiglass divider. Animals were handled for a minimum of 5 days and then used 7 to 11 days after cuff/sham surgery. Due to the relatively early experimental endpoint wound clips were not removed. Immediately following slice preparation, the right sciatic nerve was exposed, and the experimenter confirmed the cuff placement.

### Chronic constriction of the infraorbital nerve surgery

Male and female mice were anesthetized using a mixture of ketamine and xylazine (i.p) and placed in a supine position on a temperature-controlled heating pad. Using aseptic surgical techniques, the infraorbital branch of the trigeminal nerve was exposed through an intraoral incision and freed from surrounding connective tissue. Approximately 1 to 2 mm from its point of exit at the infraorbital foramen, the nerve was loosely ligated with sterile, 4-0 silk sutures. The incision was closed with VetBond tissue adhesive, and the mice were monitored continuously until fully recovered from anesthesia. Daily post-surgical monitoring continued for 5 to 7 days while the mice continued to recover in their home cage.

### von Frey testing

#### Sciatic cuff group

In a subset of animals von Frey testing was performed 7 days after surgery as previously described (Torres-Rodriguez *et al*., 2023). Male mice were habituated in the von Frey apparatus in a red-lit room for 2 hr prior to testing. The smallest filament that elicited a paw withdrawal response in at least three of five trials was taken as the mechanical threshold for each hind paw. Tactile sensitivity was determined for both hindpaws by an experimenter blind to pain treatment.

#### CCI-ION group

We confirmed the presence of CCI-induced, mechanical allodynia in subset of mice (5 sham and 5 CCI-ION, male ∼ 10 weeks old) using a 1.4 g Von Frey filament (Ugo Basil). Male and female mice were gently held in the hand of the experimenter and the filament was applied to the dorsal edge of the vibrissae pad ipsilateral to the ligated nerve. The number of nocifensive responses to 5 applications of the filament was recorded. Nocifensive responses were defined as the animal actively withdrawing its head, or aggressively biting or swiping at the filament. We tested for differences in the fraction of nocifensive responses between groups at each time point (pre-surgery/baseline and again 1 and/or 2 weeks after surgery) using a mixed effect model with Sidak’s post-hoc multiple comparison test.

### Acute slice preparation

A laboratory from one institution performed recordings from the sciatic cuff model, while a laboratory from a second institution performed recordings from the CCI-ION model. We used similar recording protocols with the exceptions noted below. **Sciatic cuff group**: Mice were deeply anesthetized with 1.2 mL of intraperitoneal Avertin (1.25%). They were then transcardially perfused with approximately 30 mL of an ice-cold, choline-based, cutting solution (110.0 mM choline chloride, 25.0 mM NaHCO_3_, 25 mM D-glucose, 12.7 mM L-ascorbic acid, 7.2 mM MgCl_2_, 3.1 mM pyruvic acid, 2.5 mM KCl, 1.25 mM NaH_2_PO_4_, 0.5 mM CaCl_2_). The brain was extracted and 300 µm coronal slices were prepared using a Leica VT1200 S vibrating blade microtome (Leica Microsystems Inc.). Slices containing the right PBN were then transferred to a room temperature recovery chamber for at least 1 hr prior to recording. The recovery chamber contained room temperature aCSF (125 mM NaCl, 25 mM D-glucose, 25 mM NaHCO_3_, 2.5 mM KCl, 2 mM CaCl_2_, 1.25 mM NaH_2_PO_4_, 1 mM MgCl_2_). Both solutions were continuously perfused with a carbogen mixture (95%/5% O_2_/CO_2_).

#### CCI-ION group

Mice were anesthetized with a mixture of ketamine and xylazine and transcardially perfused with an ice-cold, NMDG-based, cutting solution of (in mM): 92 NMDG, 30 NaHCO_3_, 20 HEPES, 25 glucose, 5 Na-ascorbate, 2 thiourea, 1.25 NaH_2_PO_4_, 2.5 KCl, 3 Na-pyruvate, 0.5 CaCl_2_ and 10 MgSO_4_. Horizontal brain sections, 300 μm thick, from both left and right PBN were collected and used for recordings.

### Whole-cell slice electrophysiology

Following recovery, slices were transferred to the recording chamber. Slices were continuously perfused (1-2 mL/min) with oxygenated aCSF. The recording chamber was maintained at 32-34°C using an inline solution heater and a heated platform under the control of a TC-344C Dual Channel Temperature Controller (sciatic cuff) or room temperature (CCI-ION). Neurons were identified using differential interference contrast on an Eclipse FN1 (Nikon) or BX-51 (Olympus) microscope. Freshly pulled glass micropipettes filled with a potassium methylsulfate/gluconate-based internal solution were used in all bridge-mode recordings: one lab (in mM): 120 CH_3_KO_4_S, 20 KCl, 14 Phosphocreatine, 10 HEPES, 8 NaCl_2_, 4 Mg-ATP, 0.3 Tris-GTP, 0.2 EGTA. Another lab (in mM): 120 K-gluconate, 10 KCl, 10 HEPES, 1 MgCl_2_, 0.5 EGTA, 2.5 Mg-ATP, 0.2 Tris-GTP. For voltage clamp recordings examining inhibitory currents, we used a high chloride intracellular solution (in mM): 70 K-Gluconate, 60 KCl, 10 HEPES, 1 MgCl_2_, 0.5 EGTA, 2.5 Mg-ATP, 0.2 GTP-Tris, and included 50 μM APC and 20 μM CNQX in the recording ACSF. Signals were acquired using Axon Multiclamp 700B (Molecular Devices) and amplified using an Axon Digidata 1550 (Molecular Devices). Data was sampled at 100.00 kHz and low-pass filtered at 10 kHz.

### Electrophysiological measurements

Capacitance and input resistance were calculated from a 25ms ±10 mV voltage step from −70 mV. Repetitive action potential firing was evoked using 500ms depolarizing current steps of increasing magnitude (5pA). This protocol was performed both on cells at their resting membrane potential (quiescent cells), or with additional current applied to maintain the cell at −70 mV (spontaneous cells). To minimize the impact of small changes in V_rest_, rheobase was defined as the minimum current required to evoke an action potential in response to a depolarizing current step (500 ms) where the cell also fired to the next step.

### Action potential analysis

In spontaneously firing neurons, the average action potential frequency was calculated from a 1 min period of stable firing without holding current. The Easy Electrophysiology software package (v2.5.0) was used to measure action potential kinetics (RRID:SCR_021190). The values from 20 action potentials were averaged for each cell. Voltage threshold was defined using a first derivative threshold of 33 mV/ms.

### Paired Pulse Ratios

All recordings of electrically evoked inhibitory postsynaptic currents (IPSCs) where performed in the presence of 20 μM CNQX and 50 μM APV to suppress AMPA and NMDA responses respectively. IPSCs were evoked by passing current through a bipolar stimulating electrode at the edge of PBN. Stimulus intensities were determined for each cell by finding the lowest current threshold which reliably evoked an IPSC. Paired stimuli within a trial were 50 ms apart with 5 to 10 seconds between trials. Paired pulse ratios were determined by averaging 10 to 20 individual trials and dividing the amplitude of the second response by the amplitude of the first response.

### Statistics

Data are represented as mean ± 95% confidence interval (Figures 1-5) or median ± 95% confidence interval (Figures 6-7). Repeated-measured 2-way ANOVA, 1-way ANOVA, mixed-effects analysis, Chi-square, student’s t-tests, Man-Whitney and Wilcoxon matched pairs test were performed as appropriate using GraphPad Prism version 9.4.0 for Windows (GraphPad Software, San Diego, California USA, www.graphpad.com). A single ROUT outlier LF capacitance was removed from the dataset.

## Results

### Neuropathic injury model increases spontaneous activity in PBN neurons

To test the hypothesis that lPBN neurons are more excitable following injury, we used the sciatic cuff model of neuropathic pain (Benbouzid et al., 2008). Animals received either cuff implantation or sham surgery to the sciatic nerve 7-11 days prior to acute brain slice recordings (**Figure 1A**). To validate that cuff implantation in the sciatic nerve induces tactile hypersensitivity, we performed the von Frey test 7 days after sciatic nerve surgery in a subset of animals. As expected, sham animals did not show hypersensitivity in the hindpaw ipsilateral to sciatic nerve surgery, compared to the contralateral hindpaw. In contrast, cuff animals presented hypersensitivity in the hindpaw ipsilateral to the sciatic cuff implantation, compared to the contralateral hindpaw (**Figure 1B**; 2-way ANOVA: interaction F(1, 13)= 54.03, p<0.0001; surgery group F(1, 13)= 31.23, p<0.0001; paw F(1, 13)= 84.25, p<0.0001; Bonferroni’s multiple comparison: sham p=0.4075; cuff p<0.0001).

**Figure 1.**
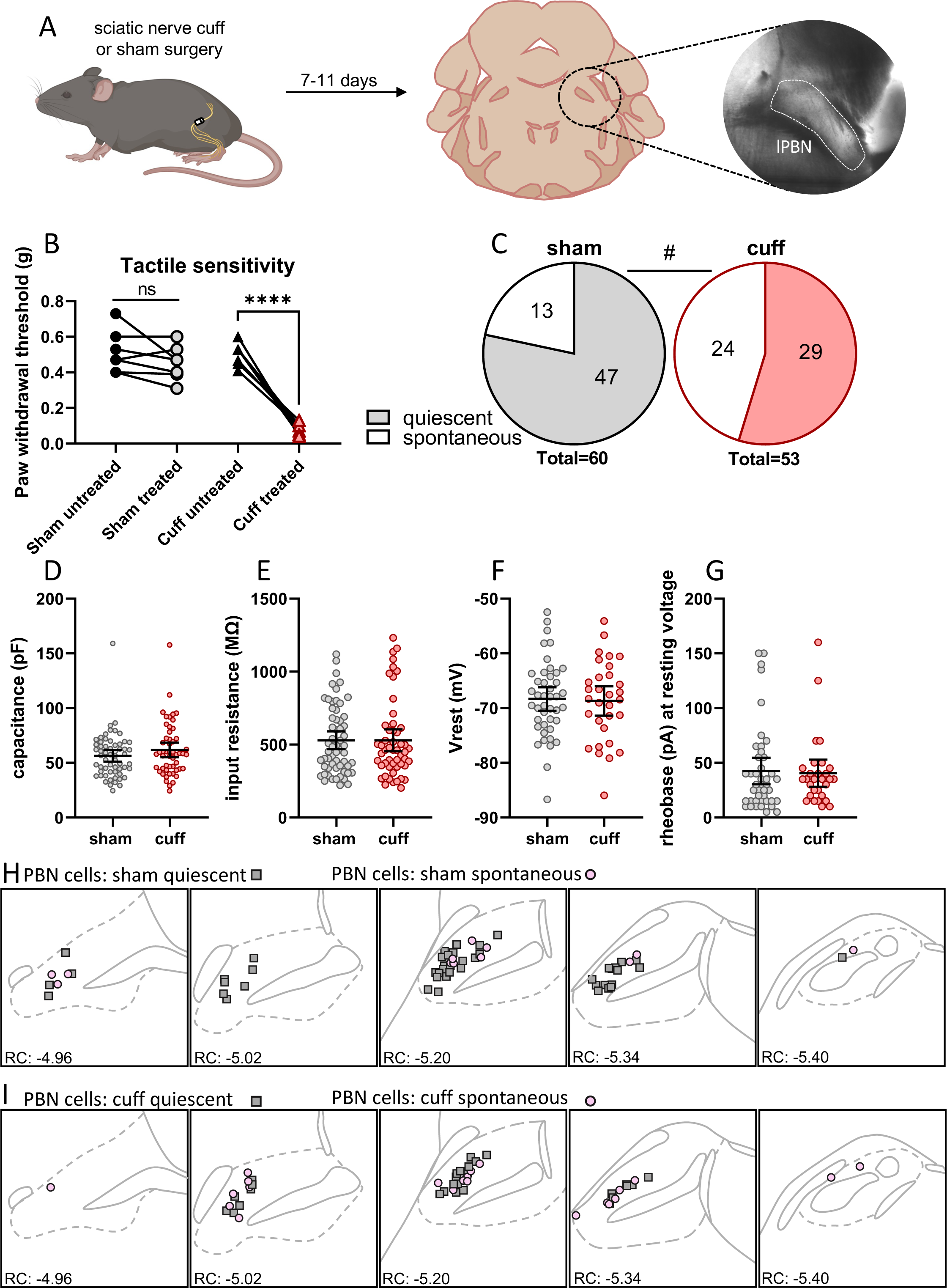
Neuropathic injury model increases spontaneous activity in PBN neurons. **(A)** Mice were used 7-11 days after sciatic cuff surgery for slice electrophysiology in pBN. **(B)** Cuff surgery induced tactile hypersensitivity, but sham surgery had no effect. **(C)** Proportion of spontaneously-firing neurons in each firing category from sham (22%, 13/60) and cuff (45%, 24/53) animals. Neuronal properties, including **(D)** capacitance, **(E)** input resistance, **(F)** resting membrane potential, **(G)** and rheobase were similar in sham and cuff animals. Anatomical location of the recorded neurons from **(H)** sham and **(I)** cuff animals. Gray squares: quiescent cells. Pink circles: spontaneously firing cells. Bonferroni corrected t-test: ****p<0.0001. χ^2^ test: #p<0.05

Using whole-cell patch-clamp electrophysiology in acute brain slices, we found that a proportion of neurons fired spontaneously (i.e. in the absence of holding current) in the sham condition (22%, 13/60 cells). We have previously shown that PBN neurons can spontaneously fire in pain-free animals *in vivo* (Uddin et al., 2018). The proportion of spontaneously active neurons recorded in cuff animals (45%, 24/53 cells) was significantly higher (**Figure 1C**; χ^2^ =6.726, df=1, z=2.597**, p=0.0095**). We examined the intrinsic properties of these neurons and found no significant differences in capacitance (**1D**) or input resistance (**1E**). Resting membrane voltage (**1F**) and rheobase (**1G**) were also not significantly different between quiescent neurons from sham and cuff animals.

To determine whether spontaneous neurons were located in any particular lPBN subnuclei, we mapped the location of the recording pipette following each recording (**Figure 1H-I**). Spontaneous neurons were intermingled with quiescent neurons and recording locations were similar between sham and cuff groups. These data show that injury-induced increases in PBN spontaneous activity observed *in vivo* (Matsumoto et al., 1996; Raver et al., 2020) are preserved in an *ex vivo* acute brain slice preparation. Our results agree with findings suggesting that injury-induced PBN hyperexcitability may underlie pain hypersensitivity (Matsumoto et al., 1996; Raver et al., 2020; Sun et al., 2020; Uddin et al., 2018).

### Spontaneous firing frequencies and action potential properties of individual neurons are unaltered in a neuropathic injury model

The increase in the number of spontaneously firing PBN neurons from cuff animals suggests that the intrinsic membrane properties, firing frequencies or single action potential waveforms of these neurons may be impacted by the cuff model (**Figure 2A-B**). We re-examined the intrinsic properties shown in **Figure 1**, restricting our analysis to neurons that were firing spontaneously. The capacitance (**Figure 2C**), input resistance (**Figure 2D**), and average firing frequency (**Figure 2E**) of spontaneously firing neurons were not significantly impacted by neuropathic injury. Examination of the single action potential waveforms in these spontaneously active neurons showed that voltage threshold (**Figure 2F**) and peak voltage (**Figure 2G**) were not significantly different between groups. The rise time from voltage threshold to peak (**Figure 2H**), decay from peak to 90% of threshold (**Figure 2I**), and action potential width (**Figure 2J**) were not significantly different between sham and cuff groups. These results demonstrate that, while the number of spontaneously-firing PBN neurons increases following neuropathic injury, this effect does not arise from changes in intrinsic properties or action potential kinetics.

**Figure 2.**
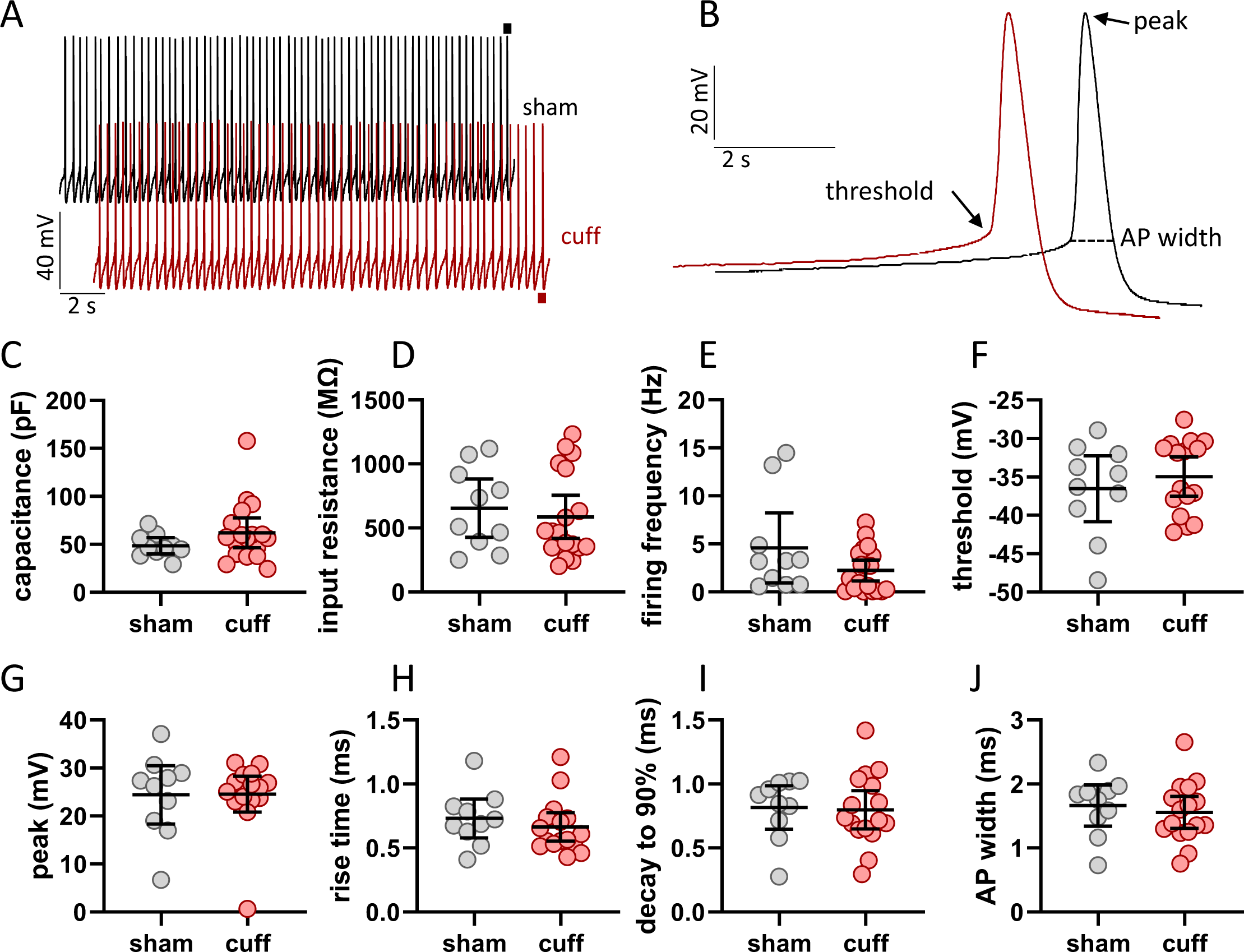
Spontaneous firing frequencies and action potential properties are unaltered in a neuropathic injury model. **(A)** Example traces of spontaneous firing in neurons from sham (black) and cuff (magenta) animals. **(B)** Example action potentials from panel A indicated by horizontal bars. There were no differences between neurons from sham and cuff cells in **(C)** capacitance, **(D)** input resistance, **(E)** firing frequency, **(F)** action potential threshold, **(G)** peak, **(H)** rise time from threshold to peak, **(I)** decay from peak to 90% of threshold or **(J)** total action potential width.

### Firing properties of PBN neurons are heterogeneous

We next examined the firing properties evoked by a prolonged (500 ms) depolarizing current step. We distinguished 4 firing types: spontaneously-firing, regular spiking (RS), late-firing (LF), and reluctant firing (**Figure 3A**). Neurons that fired an action potential within 30 ms of the beginning of the current step were classified as RS (27%, 16/60 sham cells), and those that fired after that point were classified as LF (42%, 25/60 sham cells). Both RS and LF neurons fired repetitively without much variation in inter-spike-interval throughout the series of steps. Neurons that did not fire in response to the 80pA depolarizing current injection, but fired at higher current injection steps, were classified as reluctant (8%, 5/60 sham cells). When spontaneously-firing neurons were held at −70 mV they also largely fell within these categories. 5 neurons did not fall into any of these categories and were excluded from further analysis (**Figure 3B**). Two neurons presented bust-like firing at low current steps, and 3 neurons showed strong depolarizing block.

**Figure 3.**
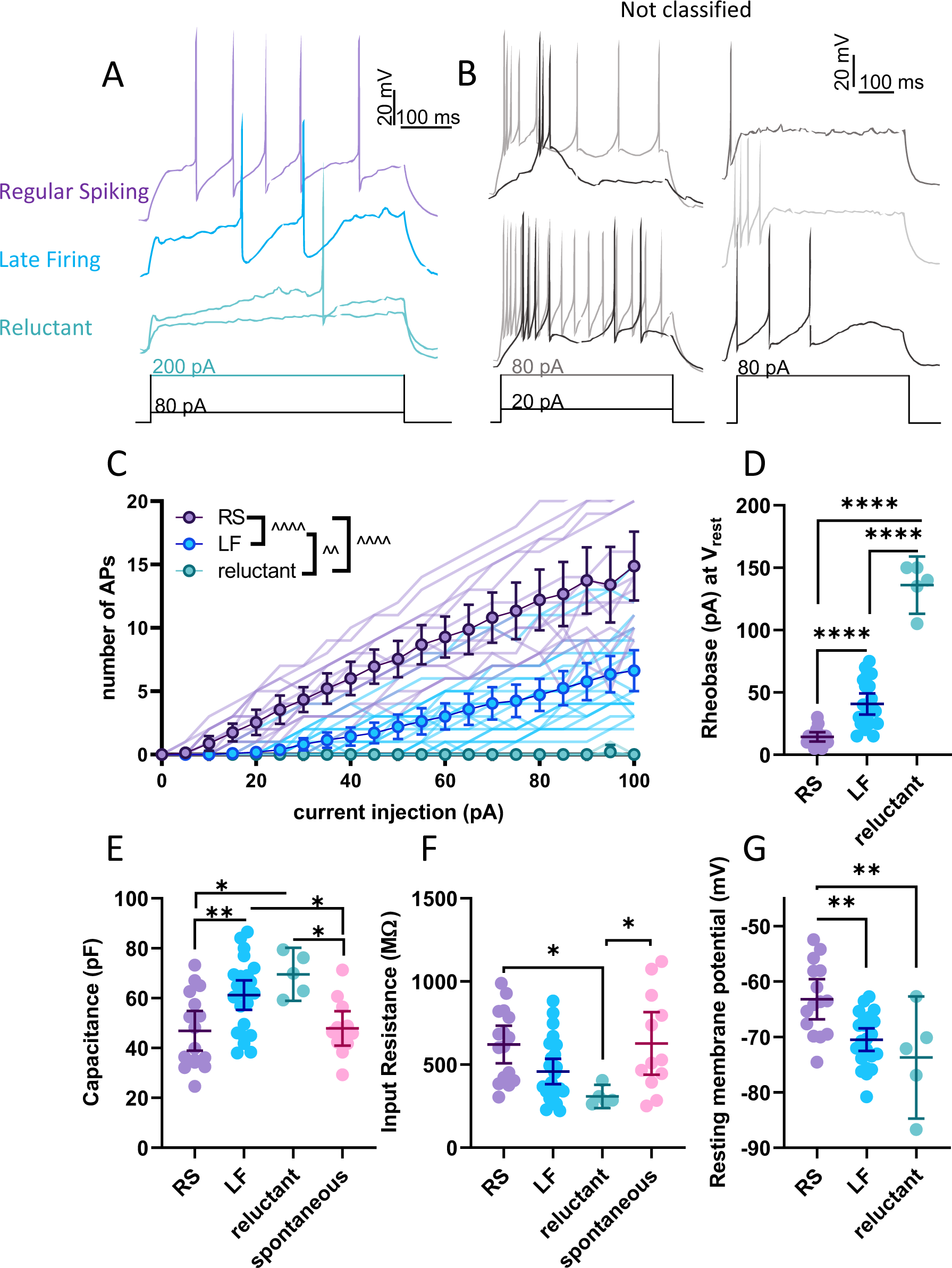
Heterogeneous firing types of PBN neurons. **(A)** Representative traces of regular spiking, late firing, and reluctant neurons. **(B)** Representative traces of neurons that did not fit the main firing types. **(C)** Input-frequency and **(D)** rheobase for each firing type. Intrinsic properties, including **(E)** capacitance, **(F)** input resistance, and **(G)** resting membrane potential were all different between firing types. Data for spontaneous neurons are replicated from Figure 2 for comparison. Each datapoint represents a cell. Bonferroni corrected t-test: *p<0.05, **p<0.01, ****p<0.0001. Main effect from mixed-effects analysis: ^^p<0.01, ^^^^p<0.0001.

To evaluate the excitability of each firing type we analyzed their evoked firing in response to a series of prolonged (500 ms) depolarizing current steps. Both RS and LF neurons fired increased number of spikes as a function of the amplitude of depolarizing current injections (**Figure 3C**). In contrast, reluctant neurons rarely fired within the amplitude steps shown here. Quantification of the input-frequency relationship between firing types showed that RS neurons were the most excitable and reluctant neurons the least. (Mixed-effects analysis: current step F(1.498, 56.83)= 68.09; firing type F(2, 38) = 37.39; interaction F(40, 759)= 23.28; all p<0.05). The differences in excitability between firing types was also reflected in the rheobase (**Figure 3D**; One-way ANOVA: F(2,40)=122.0, **p<0.0001**; all pairwise comparisons **p<0.0001**), with RS neurons showing the lowest rheobase and reluctant the highest.

Intrinsic membrane properties can dictate firing patterns and strongly influence neuronal output. To gain insight into the mechanisms underlying these heterogenous firing patterns, we examined the intrinsic properties of each firing type. PBN neurons with different firing types had significantly different capacitances (**Figure 3E**; One-way ANOVA: F(3,53)=6.815, p=**0.0006**), input resistances (**Figure 3F**; One-way ANOVA: F(3,54)=4.508, p=**0.0068**), and resting membrane potentials (**Figure 3G**; One-way ANOVA: F(2,40)=9.263, p=**0.0005**). Post hoc analyses further revealed that reluctant neurons have significantly higher capacitance values than spontaneous and RS neurons (Bonferri-corrected post hoc vs spont **p=0.0398**, vs RS **p=0.0101**), as did LF neurons (vs spont **p=0.0398**, vs RS **p=0.0096**). Reluctant neurons also had the lowest average input resistance of all firing types (vs spont p=**0.0422;** vs RS p=**0.0364**) and the most hyperpolarized average resting membrane potential. Lastly, resting membrane potential in RS neurons was significantly more depolarized than either LF or reluctant cells (vs LF p=**0.0018**; vs reluctant p=**0.0041**). Taken together, these data demonstrate that the lPBN is composed of cells with different firing types that have different intrinsic membrane properties and firing responses to depolarizing current injections.

### Neuropathic injury model bidirectionally impacts evoked firing

As most neurons fell into the RS or LF categories, we focused on those populations in evaluating the effects of neuropathic injury on intrinsic membrane properties and excitability. In RS and LF neurons, there was no difference in capacitance or input resistance between cuff and sham groups (**Table 1**). We next examined evoked firing in response to prolonged depolarizing current injections, as we hypothesized that the increased responsivity to sensory stimuli described *in vivo* (Matsumoto et al., 1996; Raver et al., 2020; Uddin et al., 2018) would translate as a magnified response to depolarizing current injection. Notably, RS neurons from cuff animals fired *fewer* APs than those from sham animals (**Figure 4A-B**; Mixed-effects analysis: current step F(1.567, 29.46) = 86.22, p<0.0001; surgery F(1, 19) = 8.090, p=0.0104; interaction F(20, 376) = 3.903, p<0.0001) without a concurrent increase in rheobase (**Figure 4C**; p=0.1221). This hypo-excitability was not due to a difference in resting membrane potential, as the resting membrane voltage was not significantly different between conditions (**Figure 4D**; p=0.3010). In LF neurons, however, there were no significant differences in evoked firing frequencies (**Figure 4E**), rheobases (**Figure 4F**) or resting membrane potentials (**Figure 4G**). These data suggest that, contrary to our hypothesis, injury reduces evoked excitability in a subset of PBN neurons.

**Figure 4.**
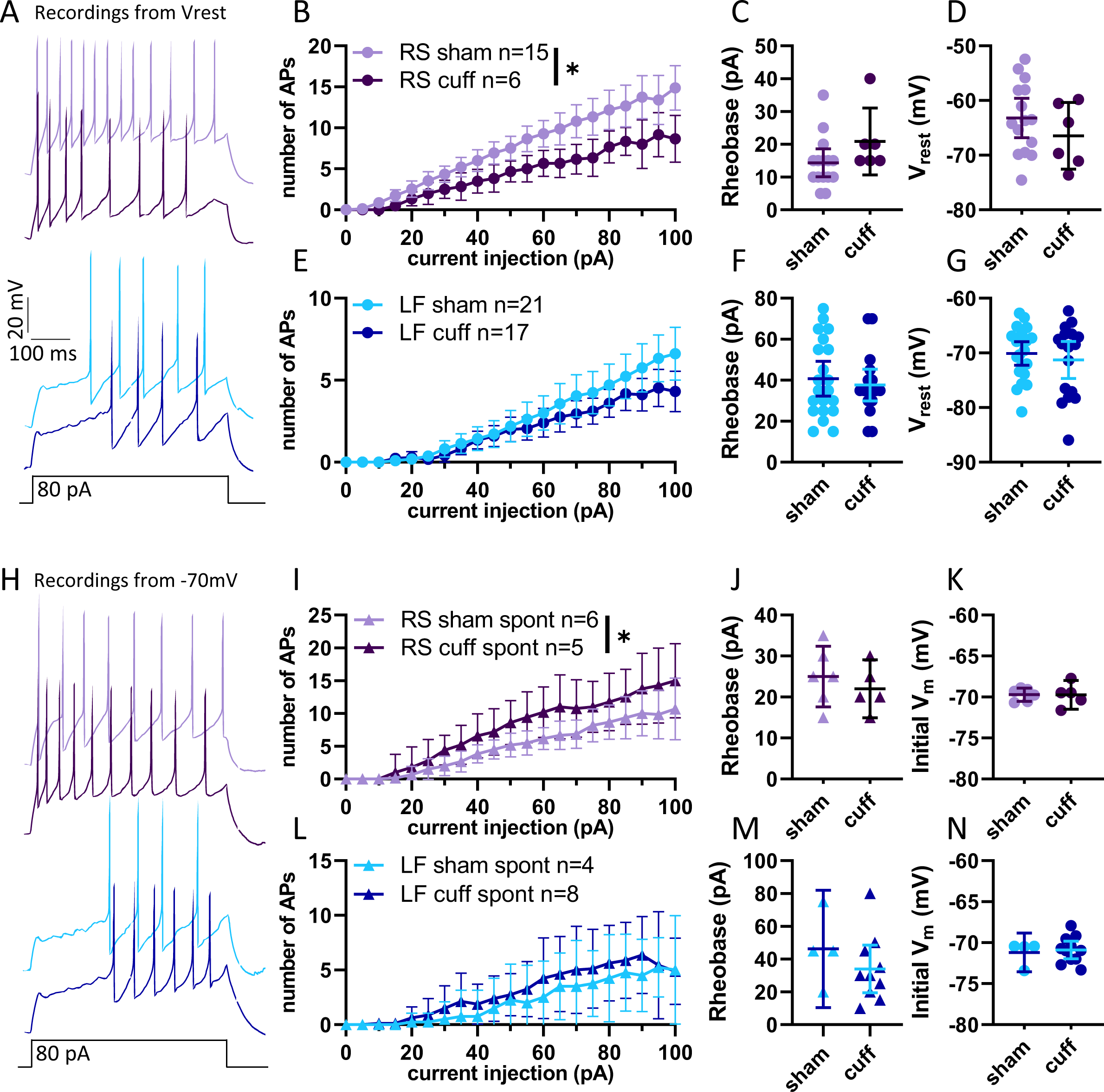
Neuropathic injury model bidirectionally impacts evoked firing. **(A)** Representative traces of action potentials in RS and LF neurons from sham and cuff animals. **(B)** Quiescent RS neurons were more excitable in sham animals than cuff animals, though **(C)** rheobase and **(D)** resting membrane potential were not different. **(E)** Quiescent LF neurons were not impacted by neuropathic injury in either repetitive firing **(F)** rheobase, or **(G)** resting membrane potential. **(H)** Representative traces of spontaneous firing in RS and LF neurons from sham and cuff animals. **(I)** Input-frequency relationship for RS neurons held at −70 mV. **(J)** Rheobase and Initial membrane voltage **(K)** spontaneous RS neurons. **(L)** Input-frequency relationship for LF neurons held at −70 mV. **(M)** Rheobase and Initial membrane voltage **(N)** spontaneous LF neurons. Data for sham neurons is replicated from figure 3 for comparison. Main effect from mixed-effects analysis: *p<0.05.

**Table 1.**

|  | Quiescent |  |  |  | Spontaneous |  |  |  |
| --- | --- | --- | --- | --- | --- | --- | --- | --- |
|  | RS |  | LF |  | RS |  | LF |  |
|  | sham | cuff | sham | cuff | sham | cuff | sham | cuff |
| Capacitance (pF) | 46.85±14.94<br>(n=16) | 55.4±21.08<br>(n=7) | 65.11±23.91<br>(n=25) | 59.84±15.89<br>(n=18) | 46.44± 12.59<br>(n=7) | 50.21±23.54<br>(n=6) | 50.05±10.32<br>(n=4) | 59.72±25.83<br>(n=11) |
| Rin (MΩ) | 620.6±212.8<br>(n=16) | 481.7±170.8<br>(n=7) | 458.0±184.0<br>(n=25) | 551.6±264.0<br>(n=18) | 581.8± 298.5<br>(n=7) | 563.3±307.5<br>(n=6) | 734.1±344.0<br>(n=4) | 590.7±302.0<br>(n=11) |
| Rheobase at native<br>V<sub>rest</sub> (pA) | 14.33±7.76<br>(n=15) | 20.83±9.704<br>(n=6) | 40.71±18.59<br>(n=21) | 37.65±15.12<br>(n=17) | N.A. | N.A. | N.A. | N.A. |
| Rheobase at -70 mv<br>(pA) | 33.08±20.37<br>(n=13) | 50.0±42.43<br>(n=7) | 42.37±29.46<br>(n=19) | 40.88±18.31<br>(n=17) | 25.00± 7.071<br>(n=6) | 21.67±5.164<br>(n=6) | 46.25±22.50<br>(n=4) | 34.00±20.39<br>(n=10) |

To examine the evoked firing properties of spontaneously firing neurons, we held them at −70mV and applied prolonged depolarizing current steps to elicit firing (**Figure 4H**). Spontaneous neurons from sham animals held at −70mV were classified as RS and LF neurons based on their firing responses to depolarizing current injections. Cuff treatment did not measurably impact capacitance or input resistance for either spontaneous RS or spontaneous LF neurons (**Table 1**). In contrast to the quiescent neurons, the spontaneous neurons from cuff animals had *higher* evoked excitability, compared to those from sham (**Figure 4I**; Mixed-effects analysis: current step: F(1.765, 20.82) = 19.44, p<0.0001; treatment: F(1,12)=0.6580, p=0.4331; Interaction: F(20,236)=0.4118, p=0.9889). There was no significant change in rheobase (**Figure 4J**) in these neurons held at −70 mV (**Figure 4K**). There was no difference in the evoked excitability (**Figure 4L**) or rheobase (**Figure 4M**) of spontaneous LF neurons held at −70 mV (**Figure 4N).** Taken together these data show that there are divergent changes in injury-induced excitability in a subset of PBN neurons with higher excitability observed in one population (spontaneous RS) and lower excitability in a second (quiescent RS).

### Parabrachial After Discharges ex vivo do not arise from changes in intrinsic properties of PBN neurons

In addition to enhanced responses to noxious stimulation, PBN neurons recorded in chronic pain models *in vivo* often continue firing at elevated levels long after the stimulus has ended (Raver et al., 2020; Uddin et al., 2018). Like the changes in evoked excitability described above, these after-discharges may contribute to the exaggerated pain behaviors observed in these models. In a second series of experiments, we sought to determine whether after-discharges are intrinsic to PBN neurons or arise from extrinsic factors, such as alterations in synaptic strength. Using a similar model of neuropathic pain, chronic constriction injury of the infraorbital nerve (CCI-ION), we used whole cell patch clamp techniques to test the prediction that after-discharges are more prevalent in PBN neurons from injured mice **(Figure 5A)**. These recordings were performed in a different laboratory than those focused on the intrinsic properties described above but using similar *ex vivo* techniques. Differences in the recording conditions are described in the Methods.

**Figure 5:**
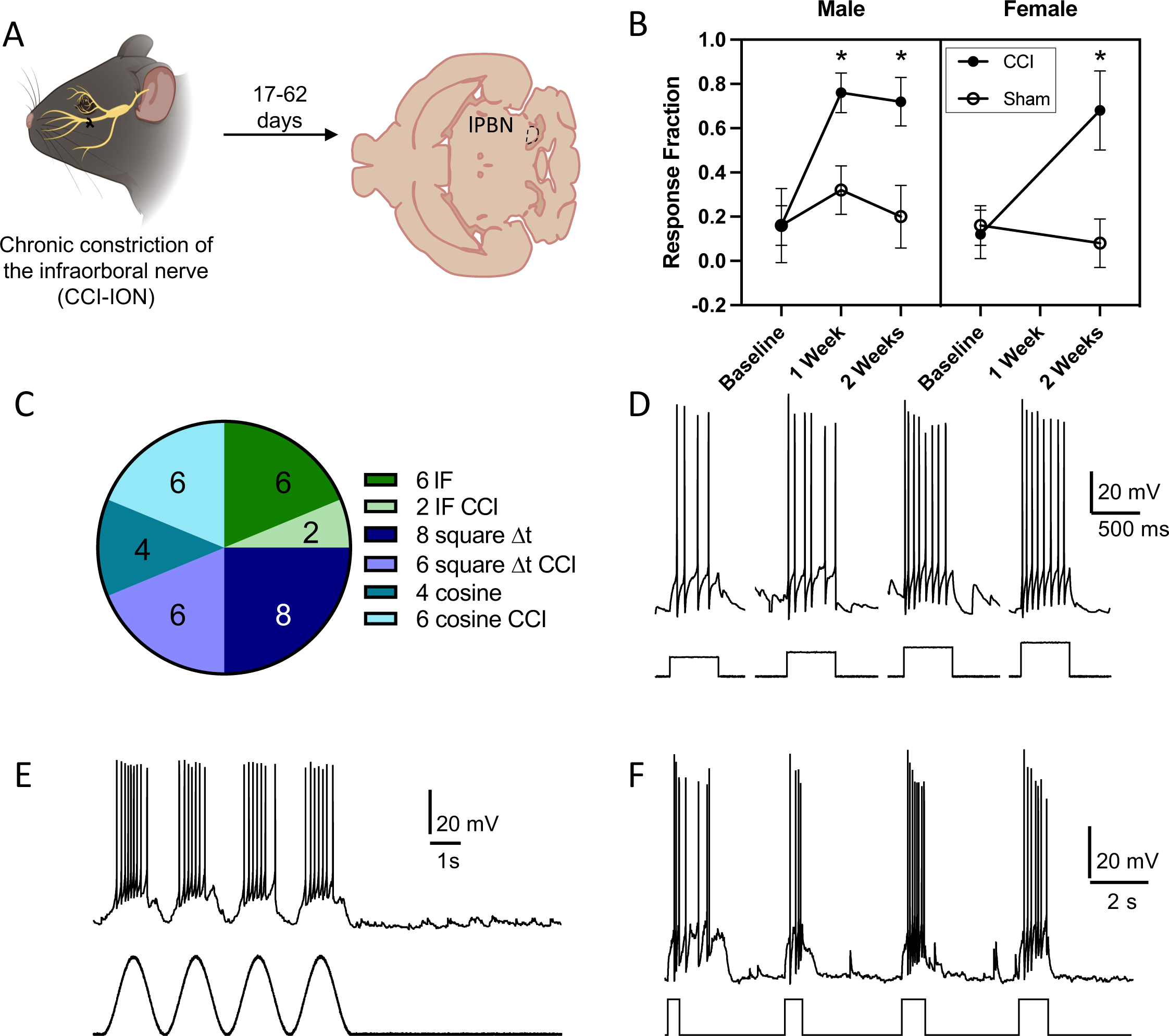
After discharges do not reflect intrinsic membrane properties of PBN neurons. **(A)** Experimental design for investigating after-discharges. CCI-ION was used as a model for peripheral nerve injury and electrophysiological data were acquired from horizontal sections through the PBN. **(B)** CCI-ION results in mechanical hypersensitivity in both sexes that persisted for at least two weeks after injury **(C)**. Summary of the number of neurons from sham and CCI mice tested with different current injection protocols in attempts to evoke after-discharges. We recorded from a total of 12 neurons from sham and 11 neurons from CCI mice, but some PBN neurons were tested with multiple protocols. Examples of each protocol include: square pulses of fixed duration and increasing magnitude **(D)**, depolarizing sinusoidal currents **(E)**, and square pulses of fixed intensity and increasing duration **(F)**. After-discharges, such as the one following the first depolarizing square pulse in F, were rarely observed, and could not be consistently evoked, in any PBN neuron. Mixed effects analysis with Sidak’s multiple comparison test, * p < 0.05

Similar to the tactile hypersensitivity observed in mice with a sciatic nerve cuff, male and female mice with a CCI-ION developed mechanical allodynia in the vibrissae pad innervated by the ligated nerve **(Figure 5B).** When measured one to two weeks after injury, mice of both sexes in the CCI-ION group showed a higher frequency of nocifensive responses to 5 normally innocuous mechanical stimuli applied to the vibrissa pad relative to shams (mixed-effects model with Sidak correction for multiple post-hoc comparisons, p < 0.0001 at one and two weeks, n = 5 per group).

We recorded from a total of 23 neurons (sham: n = 12 neurons from 1 male and 3 female mice, CCI-ION: n = 11 neurons from 4 female mice), all of which displayed one of the firing patterns described above. Most were regular spiking (RS, n = 8 sham and 7 CCI) and the remaining were either late firing (LF, n = 1 sham and 4 CCI), or fast spiking (FS, n = 3 sham). Our lower sample size in this model prevents statistical comparison of intrinsic properties. However, spontaneous firing was present in 6 of 12 neurons from sham mice (50%) and 4 of 11 neurons from CCI mice (36%). In quiescent cells, the median resting membrane potential of sham neurons was −58 mV (n = 6; 95% CIs: −67 to −49 mV) and −57 mV in neurons from CCI mice (n = 5, 95% CIs: −59 to −49 mV).

We used a variety of depolarizing current pulses that varied in shape and duration as a surrogate for synaptic input, in an attempt to evoke after-discharges. These protocols included sinusoidal pulses of variable frequency (0.5 to 10 Hz) and amplitude, square pulses of fixed duration (0.5 or 1 s) but increasing intensity (2 to 3 times the rheobase), and square pulses of fixed intensity but increasing duration (0.25 to 5 s). The pie chart in **Figure 5C** shows the number of cells in which each protocol was used in an attempt to generate after-discharges. Some cells were tested with multiple protocols.

Representative recordings of each protocol are shown in **Figures 5D-F** respectively. Regardless of the parameters, the occurrence of after-discharges following any form of depolarizing current pulse was rare and observed in only 2 of 23 PBN neurons, both of which were RS neurons from CCI mice. Even in these two cells, after-discharges occurred irregularly. For example, the cell in **Figure 5F** produced an after-discharges following the first depolarizing pulse but failed to do so on subsequent depolarizations. These results suggest that the enhanced excitability and responsiveness of PBN neurons observed in chronic pain models *in vivo* are unlikely to arise from changes in intrinsic properties of the PBN neurons.

### GABA_B_ mediated presynaptic inhibition is enhanced in PBN neurons in a neuropathic injury model

An alternative mechanism for the generation of after-discharges is a shift in the balance of excitation and inhibition within PBN. Indeed, chronic pain behaviors in mice and rats with a CCI-ION injury are causally related to diminished inhibitory drive from the CeA (Raver et al., 2020). Inhibitory afferents in PBN are regulated by several classes of presynaptic receptors (Cramer et al., 2021; Teuchmann et al., 2022), with particularly strong and consistent modulation arising from GABA_B_ receptors (Cramer et al., 2021). We investigated the potential contribution of GABA_B_ receptors in injury-induced changes in PBN excitability by measuring the amplitude of pharmacologically isolated, evoked inhibitory postsynaptic currents (eIPSC) in PBN neurons (**Figure 6A**). We placed a bipolar stimulating electrode at the edge of PBN of acute brain slices and recorded neuronal responses to paired pulse stimulation in 13 neurons from 7 sham mice (2 female and 5 male) and 20 neurons from 11 CCI mice (4 female and 7 male). The mean amplitude of the second response divided by the mean amplitude of the first response is called the paired pulse ratio (PPR) and changes in this metric reflect changes in synaptic release probability(Kim and Alger, 2001; Sanabria et al., 2004). As shown in **Figure 6B**, electrical stimulation within PB reliably produced IPSCs in neurons from both sham and CCI mice. Quantification of the first eIPSC amplitude (**Figure 6C**) and PPR (**Figure 6D**) revealed no differences between the two experimental groups (PPR: sham = 1.2, 1.0 to 1.5, CCI = 1.1, 1.0 to 1.4; mean with 95% CIs, unpaired t-test, p = 0.9).

**Figure 6:**
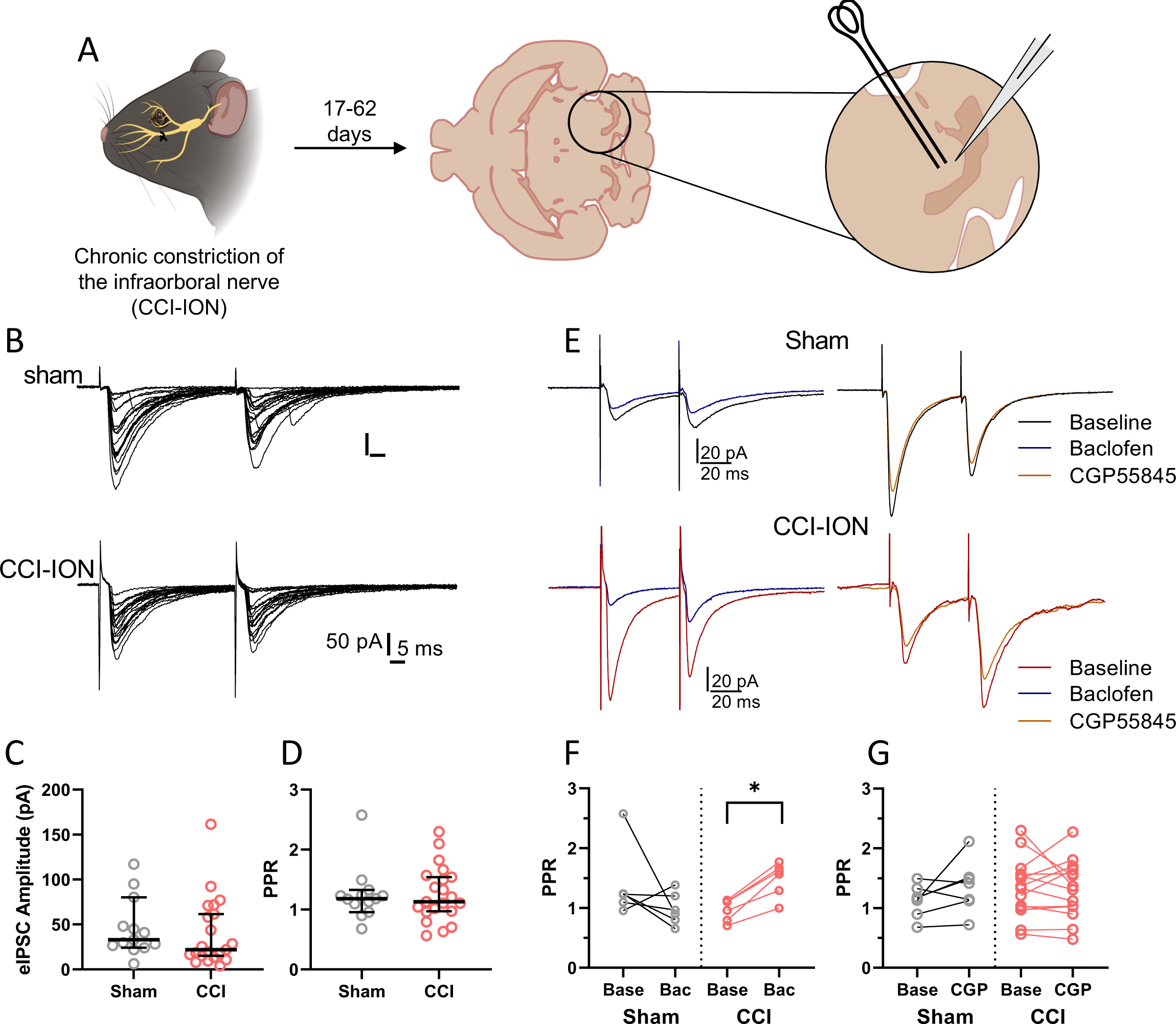
Phasic GABA_B_ inhibition of inhibitory afferents in PBN is enhanced after nerve injury. **(A)** Experimental schematic. **(B)** Samples of electrically evoked IPSCs in a PBN neuron from a sham (top) and CCI (bottom) mouse. Each depict an overlay of 20 responses to paired pulse stimulation (50 ms inter-pulse interval). All traces have been baseline corrected. Neither the amplitude **(C)** or paired pulse ratio (PPR) **(D)** was affected by CCI. **(E)** Sample recordings from sham (black) and CCI neurons (red) before and after bath application of the GABA_B_ agonist baclofen (1 μM; blue) or the GABA_B_ antagonist CGP55845 (1 μM; orange). Each trace is the average of 20 baseline-corrected individual trials. The stimulus artifact has been truncated vertically for clarity. **(E)** PPR was only affected by baclofen in PBN neurons from CCI mice. **(G)** Blocking GABA_B_ receptors did not consistently alter PPR in sham or CCI mice. Wilcoxon matched pairs test: * p < 0.05

We hypothesized that the level of tonic GABA release *ex vivo* may be insufficient to activate presynaptic GABA_B_ receptors and, consequently, any changes in the strength of this pathway may not be observable under these baseline recording conditions. To address this possibility, we measured eIPSC PPR before and after bath application of either a GABA_B_ agonist (1 μM baclofen) or antagonist (1 to 5 μM CGP-55845; **Figure 6E**). As shown in the example recordings and group data in **Figure 6F**, baclofen did not alter eIPSC PPR in PBN neurons from sham mice (baseline: 1.2, 0.8 to 2; baclofen: 0.9, 0.7 to 1.3; Wilcoxon matched pairs test: p = 0.2, n = 6). However, the median eIPSC PPR in neurons from CCI mice was significantly increased by baclofen application (baseline: 1.0, 0.8 to 1.2 95% CIs; baclofen: 1.6, 1.2 to 1.8 95% CIs; Wilcoxon matched pairs test: p = 0.03, n = 6). In contrast, the GABA_B_ receptor antagonist CGP-55845 had no effect on eIPSC PPR in neurons from either sham (baseline: 1.2, 0.9 to 1.4; CGP: 0.1.4, 1.0 to 1.8; Wilcoxon matched pairs test: p = 0.16, n = 7) or CCI mice (baseline: 1.4, 1.1 to 1.6; CGP: 1.3, 1.0 to 1.6; Wilcoxon matched pairs test: p = 0.2, n = 14, **Figure 6G**).

Together, these results indicate that, *ex vivo*, there are no differences in tonic GABA_B_ receptor-mediated suppression of inhibitory synapses in PBN following CCI. However, when activity in GABAergic afferents increases, feedback inhibition at these synapses becomes stronger in mice with CCI. This increase in GABA_B_ receptor-mediated suppression at inhibitory terminals may enable the exaggerated responses to noxious stimuli and subsequent after-discharges commonly observed this chronic pain model *in vivo*.

## Discussion

We investigated mechanisms of injury-induced PBN hyperexcitability using an *ex vivo* acute brain slice preparation. The percent of spontaneously firing neurons was higher in the cuff model of neuropathic pain, in line with *in vivo* studies demonstrating an increase in spontaneous activity in the PBN in pain. We found extensive electrophysiological heterogeneity within the lPBN of both sham and injured mice, including neurons with regular spiking (RS), fast spiking (FS) and reluctant firing patterns. Although there were no differences in the intrinsic properties within the subclasses of PBN neurons, spontaneously active RS neurons from mice with a cuff injury were more excitable compared to shams. In addition, GABA_B_-mediated presynaptic inhibition of inhibitory synapses in PBN was enhanced following nerve injury. These results suggest that increased excitability of PBN neurons, and suppression of inhibitory afferent inputs, contribute to PBN hyperexcitability and increased pain behaviors in rodent models of chronic pain.

### Neuronal heterogeneity in the PBN

Neurons in the PBN displayed significant heterogeneity in their intrinsic properties and firing patterns. Electrophysiological heterogeneity has been described previously in the lPBN, although this work has largely focused on general differences between neurons from different PBN subnuclei (Hayward and Felder, 1999; Kobashi and Bradley, 1998). Similar heterogeneity exists in gene expression, function, input targets, and output populations in the PBN (Bernard et al., 1994; Bernard and Besson, 1990; Chiang et al., 2020; Karthik et al., 2022; Pauli et al., 2022; Uddin et al., 2018; Yang et al., 2021). Although the relationships between these different functional, genetic, and anatomical properties are just beginning to be uncovered (Pauli et al., 2022), our observation that nerve injury alters the excitability of just a subset of PBN neurons suggests these factors contribute to PBN plasticity in chronic pain conditions.

### Preservation of injury-induced hyperexcitability the PBN slices

Among its many functions (Campos et al., 2018; Nagase et al., 2019; Palmiter, 2018), the PBN plays a crucial role in nociceptive and nocifensive behaviors. In response to injury, PBN neurons *in vivo* increase their spontaneous activity (Matsumoto et al., 1996; Raver et al., 2020), as well as their evoked responses (Matsumoto et al., 1996; Raver et al., 2020; Uddin et al., 2018). We find similar injury-induced changes in PBN spontaneous and activity excitability *ex vivo,* and add the novel finding that RS neurons are particularly impacted after nerve injury.

### Divergent changes in evoked excitability

Neuropathic injury induced divergent changes in the evoked excitability of RS PBN neurons (**Figure 4**). We hypothesized that PBN neurons would become hyperexcitable following injury. Consistent with this hypothesis, spontaneously firing neurons were more excitable in injured mice while quiescent neurons were *less* excitable than those in shams (Fig. 4). To our knowledge, injury-induced hypo-excitability has yet to be described in PBN, but are consistent with heterogeneity in injury-related changes in other brain regions, such as the amygdala (Li and Sheets, 2018; Wilson et al., 2019). The projection patterns of hyper- and hypoexcitable PBN neurons will be important to determine as PBN pathways that mediate different aspects of pain behaviors have been identified (Chiang et al., 2020).

### Differences between cuff and CCI-ION outcomes

In contrast to our data from the sciatic cuff model, we did not observe any differences in intrinsic excitability of PBN neurons using the CCI-ION model of neuropathic pain. This difference may arise from differences in recording conditions: 32-34°C for recordings from the sciatic cuff mice versus room temperature (∼24 °C) for CCI-ION recordings. Sex based differences in the response of PBN neurons to injury may also be a factor, since our cuff/sham data are from male mice while our data from CCI-ION are from a mix of male and female mice. Future experiments will examine these possibilities more closely.

### The role of GABA_B_ in PBN injury-induced plasticity

We did not observe significant injury-induced changes in the capacitance, input resistance, rheobase or single action potential waveforms of lPBN neurons, suggesting that alterations in intrinsic membrane properties are not a major contributor to the observed differences in excitability. Prior studies, however, suggest that deficits in GABAergic signaling may have a greater impact on PBN excitability. Long-range GABAergic inputs from the CeA are suppressed in the chronic constriction injury of the reverses pain behaviors (Raver et al., 2020). Similarly, optogenetic inhibition of GABAergic neurons in the PBN can induce pain-like behavior in uninjured mice (Sun et al., 2020). Our results suggest that changes in GABA_B_ autoinhibition may underlie these phenomena and make a larger contribution to increased PBN excitability. We found no differences in the amplitudes of electrically evoked inhibitory postsynaptic currents (IPSCs) in PBN slices from sham and CCI-ION mice. There were also no differences in IPSC paired pulse ratios (PPR) between the two groups, suggesting that the probability of synaptic release at inhibitory synapses is not affect by injury. However, addition of a low concentration of baclofen, a GABA_B_ agonist, selectively increased the IPSC PPR in slices from CCI-ION mice (Figure 6). This suggest that autoinhibition of GABAergic afferents is enhanced following nerve injury, and the resulting decrease in inhibitory drive to PBN may contribute to the hyperexcitability observed in these neurons in models of chronic pain.

### PBN circuitry: outputs and intra-PBN signaling

Specific PBN outputs have been linked to particular aspects of aversion and nocifensive responses. These include projections from PBN to the CeA (Allen et al., 2023; Chen et al., 2018; Chiang et al., 2020; Liu et al., 2022; Torres-Rodriguez et al., 2023), bed nucleus of the stria terminalis (Chen et al., 2018; Chiang et al., 2020), ventral tegmental area (Yang et al., 2021), reticular formation (Barik et al., 2018), periaqueductal gray (Chiang et al., 2020), substantia innominata (Bowen et al., 2020), and various hypothalamic (Bowen et al., 2020; Chiang et al., 2020) and thalamic nuclei (Bowen et al., 2020; Deng et al., 2020). One possible consequence of increased lPBN excitability could be increased output onto these targets, leading to increased nociception and nocifensive behaviors. Shifts onto higher firing frequencies might also have the consequence of promoting release of neuropeptide-containing dense core vesicles (Park and Kim, 2009). Calcitonin gene-related peptide (CGRP) in particular plays an important role in injury-induced plasticity in the PBN→CeA pathway (Allen et al., 2023; Han et al., 2010, 2005; Okutsu et al., 2017; Shinohara et al., 2017), and increased CGRP release have been shown to have stark consequences for CeA activity (Han et al. 2005; Shinohara et al, 2017; Okutsu et al 2017; Presto and Neugebauer 2022; Allen et al. 2023).

Hyperexcitability of PBN neurons is frequently observed in models of chronic pain and associated with exaggerated pain behaviors. Our study finds that changes in the intrinsic properties of PBN neurons are unlikely to be the primary mechanism underlying this hyperexcitability. Although subsets of regular spiking PBN neurons show injury-induced changes in excitability, our data suggest enhanced suppression of inhibitory afferents by GABA_B_ receptors make a larger contribution to the amplified responses of PBN neurons in animal models of chronic pain.

